# Establishment of the Saudi Bank of Induced Pluripotent Stem Cells (SBiPSCs): A National Platform for iPSC-Based Research and Therapy

**DOI:** 10.64898/2026.01.22.700942

**Authors:** Manal Hosawi, Moayad Baadhaim, Mohammad Al-Shehri, Gabriel Herrera-López, Gustavo Ramirez, Yara Fadaili, Samer Zakri, Ali Haneef, Fahad Hakami, Doaa Alamoudi, Nuwayir Alhusayni, Linah Aljahdali, Latifa Aljuid, Afaf Magbouli, Heba Alkhatabi, Seraj Makkawi, Ahmad Attar, Dunia Jawdat, Ahmad Alaskar, David Gomez-Cabrero, Pierre Magistretti, Jesper Tegner, Maryam Alowaysi, Khaled Alsayegh

## Abstract

**Background:** The global landscape of induced pluripotent stem cell (iPSC) resources remains heavily skewed toward European, North American, and East Asian populations, leaving the Middle East and North Africa (MENA) region critically underrepresented. This disparity hinders the application of precision medicine in populations with unique genetic backgrounds, particularly those with high rates of consanguinity and distinctive rare disease profiles. To address this gap, we established the Saudi Bank of Induced Pluripotent Stem Cells (SBiPSCs), at King Abdullah International Medical Research Centre (KAIMRC). The bank comprises two major complementary arms: one dedicated to the derivation and biobanking of iPSCs from individuals with rare and common genetic disorders, and a second focused on Human Leukocyte Antigen (HLA)-based iPSC banking to support the development of immunocompatible cell therapies.

**Methods:** SBiPSCs operates within King Abdulaziz Medical City in Jeddah under the Ministry of National Guard for Health Affairs (MNGHA)’s ethical and clinical framework. To establish the repository, we implemented a clinic-guided enrolment strategy in which treating physicians, briefed on the bank’s objectives, recruited patients with confirmed genetic diagnoses. Peripheral blood samples were collected, processed, and cells were reprogrammed using non-integrating episomal plasmids. All derived lines underwent rigorous quality control in accordance with International Society for Stem Cell Research (ISSCR) standards, including assessment of pluripotency markers, genomic integrity, and trilineage differentiation potential. To demonstrate our iPS characterization workflow and translational utility, iPSCs from a Saudi patient with familial Long QT Syndrome (LQTS) and a healthy sibling were differentiated into functional cardiac organoids. Simultaneously, for the HLA-based banking arm, the Saudi Stem Cell Donor Registry (SSCDR) database was leveraged to identify donors predicted to provide maximal coverage for the Saudi population.

**Results:** To date, SBiPSCs has successfully generated 37 iPSC lines derived from 19 Saudi patients and healthy donors. All lines exhibit robust expression of pluripotency markers, maintain normal karyotypes, and demonstrate differentiation capacity. To demonstrate our characterization pipeline and translational utility, iPSCs from an LQTS patient and a healthy sibling were generated, validated, and differentiated into beating cardiac organoids that recapitulated the disease phenotype, with microelectrode array analysis confirming prolonged field potential durations mirroring the clinical QT prolongation. Furthermore, the HLA-based banking arm has expanded to include two homozygous iPSC lines, which together provide immunological compatibility for approximately 9% of the Saudi population.

**Conclusions:** SBiPSCs represents the first centralized iPSC repository in the MENA region. The SBiPSCs is well-positioned to accelerate the translation of stem cell research into scalable, immunocompatible cell therapies and precision medicine applications aligned with national and regional healthcare priorities.

## INTRODUCTION

The full realization of precision medicine is contingent upon the availability of genomic resources that reflect the broad spectrum of human genetic variation (1,2). However, a critical “genomic divide” persists in the global landscape of available resources (3,4). Advanced research repositories and databases driving this revolution using induced pluripotent stem cell (iPSC) technologies are currently disproportionately derived from European, North American, and East Asian populations (5). This disparity leaves significant demographics, specifically the Middle East and North Africa (MENA) region, underrepresented in global stem cell biobanks (2,4,5). Consequently, the application of emerging disease models and pharmacological screenings to these populations remains limited. To bridge this gap, in 2023, we established the Saudi Bank of Induced Pluripotent Stem Cells (SBiPSCs). At the time of writing this manuscript, the SBiPSCs has successfully generated 37 characterized iPSC lines from 19 donors. The bank is poised to provide a novel resource to interrogate the molecular mechanisms of diseases in a previously unstudied genetic context.

Since their initial description by Takahashi and Yamanaka, iPSCs have become indispensable tools for modeling disease pathogenesis, high-throughput drug screening, and regenerative medicine (6–8). To democratize access to these tools and catalyze precise, mutation-guided clinical translation, major international initiatives have established comprehensive biobanks (Table 1; Fig. 1D) (9,10). In Asia, the RIKEN BioResource Research Center (BRC) and the Japanese Cancer Research Resources Bank (JCRB) serve as vital repositories cataloging thousands of lines (11,12). Similarly, the European Bank for induced Pluripotent Stem Cells (EBiSC) was established to centralize high-quality iPSCs specifically to support pharmaceutical research and drug screening (13). In North America, the New York Stem Cell Foundation (NYSCF) repository has pioneered automated derivation to create diverse ethnic panels for population-scale genetic studies (14). These consortia have been instrumental in driving disease modeling and drug discovery by providing the global research community with standardized, reliable cellular tools. However, these consortia contain very limited representation of the unique genetic architecture of the Arabian Peninsula.

**Figure 1.**
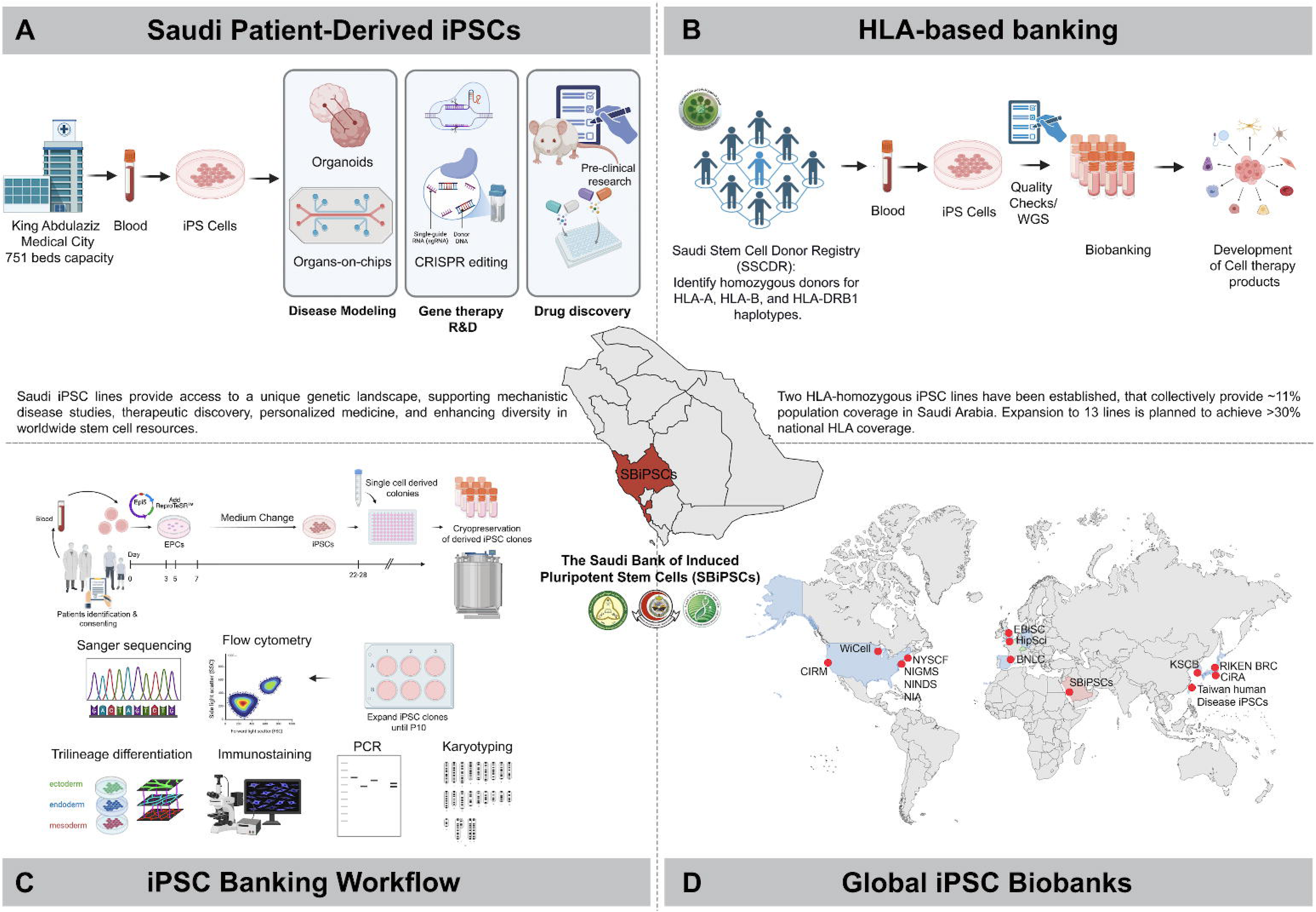
Strategic framework and operational workflow of the Saudi Bank Induced Pluripotent Stem Cells (SBiPSCs). (A) Saudi patient-derived iPSCs arm. Patients with genetic disorders are recruited from King Abdulaziz Medical City, a tertiary hospital with 751 bed capacity. Blood samples are processed and reprogrammed into iPSCs to advance disease modelling (e.g., organoids) and drug discovery capacities. (B) The HLA-based banking arm workflow. Donors are identified via the SSCDR based on homozygous HLA-A, -B, and -DRB1 haplotypes. Selected donors provide blood samples for the derivation of immunocompatible iPSC lines to develop future cell therapies. (C) The standardized reprogramming and characterization pipeline. Peripheral blood cells are reprogrammed using non-integrating episomal vectors. Resulting clones undergo rigorous quality control, including Sanger sequencing, flow cytometry for pluripotency markers, trilineage differentiation, and G-band karyotyping to ensure genomic integrity. (D) Global distribution of major iPSC repositories. The concentration of established key biobanks in North America, Europe, and East Asia, highlights the scarcity of resources in the MENA region, a gap that SBiPSCs aims to address.

**Table 1.** Summary of key international iPSC banks/repositories, their regional distribution, and demographic coverage.

| *Biobank/Repository* | *Region/Country* | *Affiliation* | *Number of Lines (Approx.)* | *Primary Focus* | *Ethnicity Represented* |
| --- | --- | --- | --- | --- | --- |
| ***WiCell Research Institute*** | North America (USA) | University of Wisconsin | 1455 iPS lines | Standardization & federal repository (global reference hESC lines; manages NHLBI “Next Gen” iPSC cohort)[3] | Primarily Caucasian, African and Hispanic |
| ***NIGMS Human Genetic Cell Repository (Coriell)*** | North America (USA) | Coriell Institute for Medical Research | 180 iPS lines | Genetic disorders & chromosomal abnormalities | Predominantly European ancestry |
| ***NIA Aging Cell Repository (Coriell)*** | North America (USA) | Coriell Institute for Medical Research | 50 iPS lines | Aging research (extreme longevity & progeria patient lines) | Predominantly European ancestry |
| ***NINDS Human Cell & Data Repository (Coriell)*** | North America (USA) | Coriell Institute for Medical Research | 193 iPS lines | Neurodegenerative diseases | Predominantly European ancestry |
| ***New York Stem Cell Foundation (NYSCF) Repository*** | North America (USA) | The Jackson Laboratory | >600 iPS lines at launch | Patient and healthy control iPS lines | A primary focus is on genetic diversity; includes an "Ethnic Diversity iPSC Panel". |
| ***CIRM iPSC Repository (Legacy)*** | North America (USA) | State Initiative (now managed by Fujifilm CDI) | ~2,500 iPS lines | Patient and healthy control iPS lines | Hispanic, African American, East Asian, and Caucasian donors (mirrors California demographics) |
| ***EBiSC (European Bank for iPSCs)*** | Europe (EU/UK) | Non-Profit Consortium | >900 iPS lines | Standardization for pharma – centralized EU iPSC bank for drug screening | Northern European (UK/EU donor base) |
| ***HipSci (Human iPSC Initiative)*** | Europe (UK) | Research Initiative | >800 iPS lines | Patient and healthy control iPS lines | White British / European (mostly UK Biobank volunteer donors) |
| ***BNLC (Spanish National Stem Cell Bank)*** | Europe (Spain) | National Node | ~ 84 iPS lines | Regional disease focus | Southern European / Hispanic (Iberian populations) |
| ***CiRA (Center for iPSC Research & Application)*** | Asia (Japan) | Public Interest Foundation | ~27 lines (clinical-grade)[34] | Clinical-grade iPSCs – HLA-homozygous | Japanese (East Asian); homozygous HLA types cover ~40% of Japan's population |
| ***RIKEN BRC (BioResource Center)*** | Asia (Japan) | National Bioresource Project | >4,200 iPS lines | Patient and healthy control iPS lines | Japanese (East Asian) |
| ***KSCB (Korea National Stem Cell Bank)*** | Asia (South Korea) | National Institute of Health | ~130 iPS lines | Clinical-grade iPSCs – HLA-homozygous | Korean (major HLA haplotypes covering >50% of S. Korea's population) |
| ***Taiwan Human Disease iPSC Consortium*** | Asia (Taiwan) | National Consortium | ~100 iPS lines | Patient and healthy control iPS lines | Han Chinese and Indigenous Taiwanese |

This “genomic gap” is particularly detrimental for Saudi Arabia and the Gulf Cooperation Council (GCC) countries, where unique population structures characterized by high rates of consanguinity have led to distinct genetic profiles and an elevated prevalence of rare and recessive disorders (15,16). The lack of regionally representative iPSC lines limits the applicability of global drug screening in Middle Eastern populations and hinders the study of locally prevalent genetic variants. Therefore, establishing the Saudi Bank of Induced Pluripotent Stem Cells (SBiPSCs) is a regional necessity and an important step toward diversifying global stem cell resources. Such a regional repository would also improve the accuracy of disease modelling and support the development of precision therapies tailored to the region’s populations.

To demonstrate the rigor of our characterization pipeline and the SBiPSC’s translational utility, we selected iPSC lines derived from a patient with Long QT Syndrome (LQTS) for in-depth validation and functional analysis. LQT Syndrome is a cardiac channelopathy characterized by delayed repolarization and a prolonged QT interval on electrocardiograms (ECG), leading to susceptibility to life-threatening arrhythmias (17,18). While the pathogenic role of *KCNH2* mutations is established, the precise molecular mechanisms driving phenotypic variability, particularly the influence of ethnic-specific genetic modifiers, remain incompletely characterized (17,19). Current heterologous expression systems often fail to capture these complex genomic interactions. Given the prevalence of cardiac channelopathies in the Saudi population and the disease’s clear electrophysiological phenotype, this model showcases the SBiPSCs’ ability to establish high-fidelity disease models that recapitulate complex clinical phenotypes in vitro. By generating functional cardiac organoids from patients with confirmed *KCNH2* mutations, we demonstrate the utility of the bank’s lines for advanced downstream applications.

In parallel, SBiPSCs is building an HLA-based iPSC repository guided by haplotype frequencies within the Saudi population. In a recent study (20), we estimated that 13 strategically selected HLA-homozygous lines could provide immunological compatibility for over 30% of the Saudi population, while 39 lines could extend coverage to more than 50%. Here, we describe the continuous expansion of this arm, which has successfully recruited two homozygous donors and generated four validated iPS clones, collectively achieving approximately 9% population coverage. In this report, we detail the establishment of the SBiPSCs at King Abdullah International Medical Research Center (KAIMRC), outlining the optimized workflows for donor recruitment and reprogramming, and discussing the potential impact of this national initiative on advancing stem-cell-based translational research across the Middle East and beyond (Fig. 1).

## Methods

### 1. Ethical Oversight

This study was conducted in accordance with the Declaration of Helsinki and was approved by the Institutional Review Board (IRB) and Research Ethics Committee of the Ministry of National Guard for Health Affairs (Protocol# NRJ22J/060/03). Adult participants provided written informed consent forms that permit the derivation of iPSCs for broad research use. In cases involving minors, consent was obtained from a parent or a legal guardian following a detailed explanation by both the primary care physician and the SBiPSCs team. All samples were de-identified and assigned a unique code to protect donor privacy.

### 2. Donor Selection and Recruitment

Two donor cohorts are actively being expanded: (1) patients with confirmed genetic disorders and (2) healthy individuals identified through the Saudi Stem Cell Donor Registry (SSCDR).

- **Patient-derived iPSCs:** Eligible participants included patients with rare or common genetic conditions verified by genetic testing (e.g., targeted sequencing or whole-exome sequencing). Individuals deemed unsuitable based on clinical judgment or lacking a confirmed genetic diagnosis were excluded. Recruitment was facilitated through collaborations with the Neuroscience and Trauma Center and King Abdullah Specialist Children’s Hospital (KASCH) at King Abdulaziz Medical City in Jeddah. Physicians, informed about the goals of the iPSC bank, recommended patients with notable genetic findings; SBiPSCs staff then obtained informed consent and scheduled blood sample collection.
- **HLA-based iPSC Banking:** Selection criteria for healthy donors followed previously established protocols (20). Individuals were included if they (i) were registered with the SSCDR, (ii) were ≥18 years of age, (iii) exhibited homozygosity at HLA-A, -B, and -DRB1 loci. Participants who have tested positive for hepatitis B or C, HIV, HTLV-1, parvovirus B19, or Treponema pallidum were excluded, as were those with any bacterial or protozoal infections or malignancies. Women who were pregnant or lactating were likewise excluded. All prospective donors considered unsuitable by the principal investigator were also excluded.

No financial incentives were provided for participation. Signed consent forms are stored by SBiPSCs staff at KAIMRC, linked only to a de-identified sample label for future reference.

### 3. Sample Collection and Processing

#### Saudi Patients-Derived iPSCs

Peripheral blood (1–3 mL) was collected in EDTA tubes during routine clinical phlebotomy. Samples were transported on ice to the KAIMRC laboratory and processed on the same day to isolate peripheral blood mononuclear cells (PBMCs) by density gradient centrifugation using SepMate^tm^ tubes (Stem Cell Technologies). Following the separation, PBMCs were either reprogrammed on the same day or stored in 90% Fetal Bovine Serum (FBS) and 10% DMSO in LN^2^ for future batch reprogramming. After thawing, frozen cells were cultured in 90% RPMI medium supplemented with 10% FBS at 37 °C with 5% CO^2^ for 24 hours before reprogramming.

#### For HLA-Based Banking

To generate iPSCs for the HLA-banking arm, Erythroid Progenitors Cells (EPCs) were used due to their genomic stability and their ability to give rise to iPSCs with a lower mutational burden compared to lines derived from other somatic cell types (21,22). Peripheral blood samples were collected in EDTA tubes and processed using the RosetteSep™ Human Progenitor Cell Basic Pre-Enrichment cocktail (Stem Cell Technologies; Cat#15226) following the manufacturer’s protocol. Isolated PBMCs were cultured for eight days in StemSpan™ SFEM II medium (Stem Cell Technologies; Cat#09605) containing 1X StemSpan™ Erythroid Expansion Supplement (Stem Cell Technologies; Cat#02692) to enrich EPCs.

### 4. Cell Reprogramming Protocol

iPSCs were generated using episomal plasmids encoding key pluripotency factors. PBMCs or EPCs (5×10^5^ cells) were electroporated using the NEON electroporation system (Thermo Fisher Scientific), carrying plasmids pCE-hOCT4, pCE-hSK (SOX2 and KLF4), pCE-hUL (L-MYC and LIN28A), pCE-mP53DD, and pCXB-EBNA1. Cells were transfected with 1 μg of each episomal vector via the Neon System. The transfected cells were then cultured on Matrigel-coated dishes and maintained in StemSpan™ medium for four days. On day four, cells received partial mTeSR medium followed by daily 50% medium exchanges until day 11 when the medium transitioned completely to mTeSR (STEMCELL Technologies). Colonies displaying characteristic iPSC morphology with tightly packed cells with a high nucleus-to-cytoplasm ratio, typically emerged around day 14. Two to three colonies per donor were manually picked, expanded, and carried forward for further validation.

### 5. iPSC Characterization and Quality Control

All iPSC lines were systematically evaluated for pluripotency, genomic integrity, and differentiation potential:

- **Immunocytochemistry:** Cells were fixed with 4% paraformaldehyde for 15 min, permeabilized with 0.1% Triton X-100 in PBS for 5 min, and blocked with 1% gelatin in PBS for 45 min. Then, cells were incubated overnight at 4 °C with primary antibodies (e.g., OCT4, NANOG and SOX2, Thermo Fisher Scientific). Appropriate secondary antibodies (Thermo Fisher Scientific) were applied for 1 h at room temperature, and nuclei were counterstained with DAPI (Table 2).
- **Quantitative reverse transcription PCR (RT-qPCR):** Total RNA was isolated using the RNeasy Kit (Qiagen) and reverse-transcribed with the High-Capacity cDNA Reverse Transcription Kit (Applied Biosystems). Pluripotency gene expression was quantified using FastStart SYBR Green Master Mix (Roche) following established protocols (23,24)
- **In vitro Trilineage Differentiation:** Stemness was further validated by differentiation into ectoderm, mesoderm, and endoderm using the STEMdiff™ Trilineage Differentiation Kit (StemCell Technologies). The confirmation of the differentiation was performed by RT-qPCR and immunocytochemistry (Table 2).
- **Flow Cytometry:** IPSCs were stained with antibodies against OCT4, NANOG, and SOX2 using the BD IntraSureTM Kit (BD Biosciences). Following the manufacturer’s instructions, reagent A was used to fix (4 × 10^5^) cells for 10 min. Then, the primary antibodies were diluted with reagent B and incubated with the cells for 30 min. Later, cells were incubated for 30 min at room temperature with secondary antibodies. The samples were analyzed using the BD FACS ARIA cell sorter.
- **Karyotyping:** Metaphase spreads were prepared following a 15 min exposure to KaryoMAX™ Colcemid™, hypotonic treatment, and fixation in methanol: acetic acid (3:1). G-banded analysis of at least 50 metaphases was performed in a clinically accredited diagnostic laboratory (High Quality Laboratories). Lines exhibiting any abnormal karyotypes were excluded from the bank.
- **Plasmid screening:** DNA was extracted using the DNA/RNA/Mini Kit (Qiagen), following the manufacturer’s instructions. During the PCR procedure, EBNA-1 primers were utilized to detect the five episomal plasmids (Thermo Fisher Scientific).
- **Short tandem repeat (STR):** Genomic DNA was extracted from iPSCs and PBMCs, then twenty-one STR loci and Amelogenin were amplified using the GlobalFiler™ PCR Amplification Kit (Thermo Fisher Scientific). The PCR amplicons were analyzed with a SeqStudio™ Genetic Analyzer. Data was collected and assessed using GeneMapper ID-X Software, performed by the Department of Pathology and Laboratory Medicine (Ministry of the National Guard–Health Affairs).
- **Mycoplasma detection:** Rapid mycoplasma detection kit was used to detect mycoplasma contamination, following the manufacturer’s instructions (AssayGenie).
- **Molecular analysis of blood-borne pathogens:** Molecular analysis of genomic DNA and RNA extracts was performed on the generated iPSC cell lines to test for Human Immunodeficiency Virus (HIV), Hepatitis B Virus (HBV), and Hepatitis C Virus (HCV) using validated molecular diagnostic assays.
- **Statistical analysis:** All statistical analyses in the study were presented as mean ± standard deviation (SD). The unpaired two-tailed Student’s *t*-test evaluated differences between samples. The *p*-value ≤ 0.05 determined statistical significance for each experimental result.

**Table 2.**
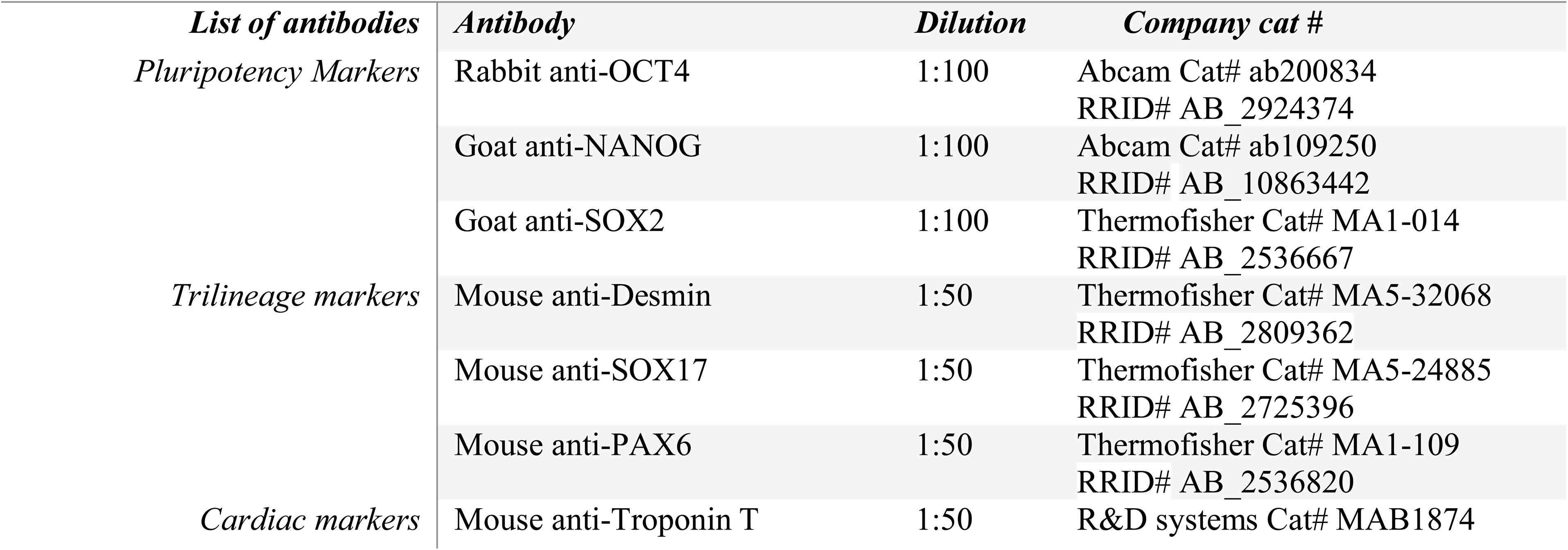
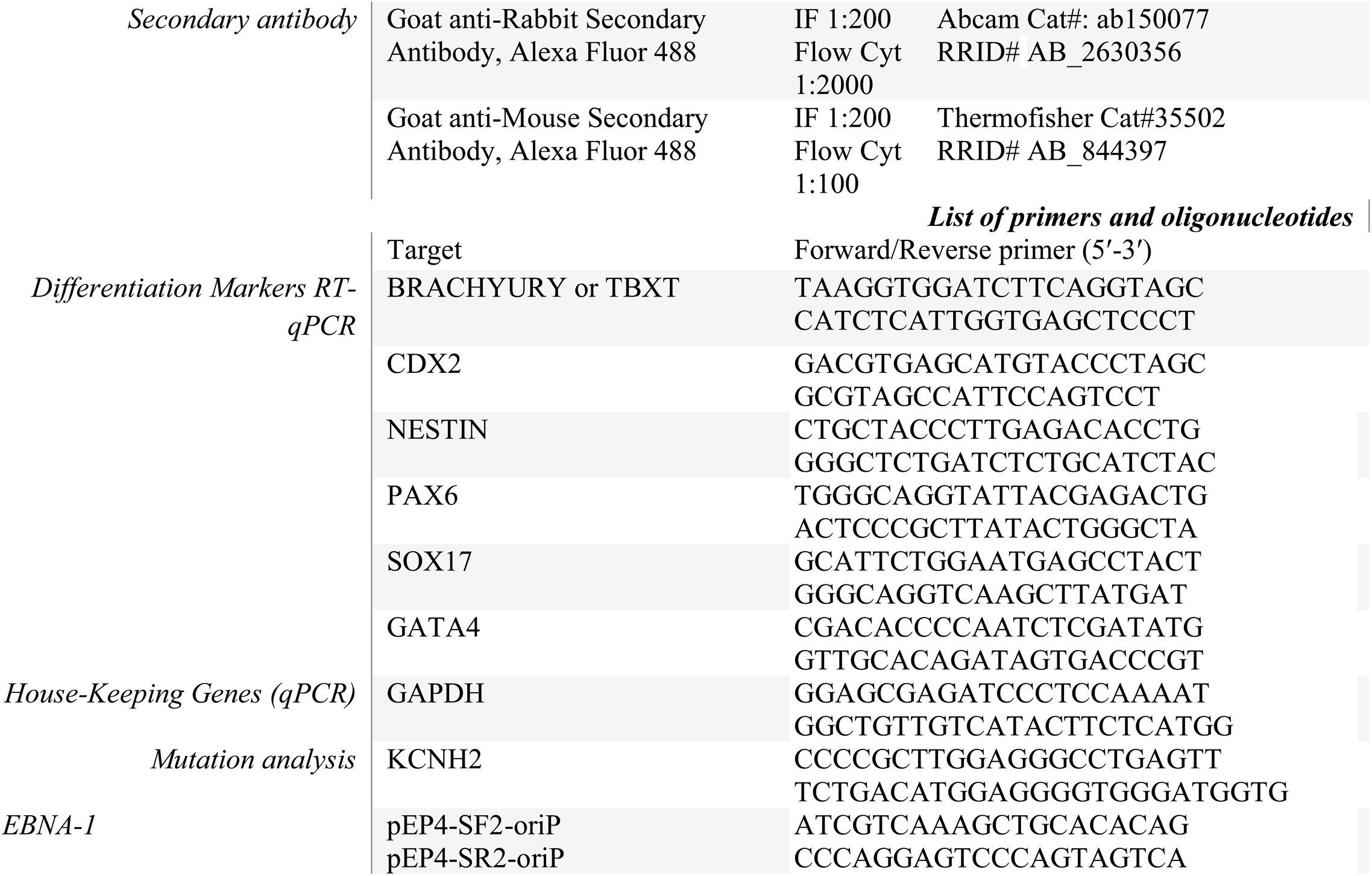
Summary of antibodies and primers used in the study.

### 6. Procedures for 3D generation and evaluation of cardiac organoids

Cardiac organoids for the LQTS-iPSC lines were generated by seeding 100,000 cells/mL in ultra-low adhesion round-bottom 96-well plates using mTeSR medium with ROCK inhibitor for 24 hours as previously described (25) (Fig. 5A). The next day, 50 μL of old medium was removed and 200 μL fresh mTeSR medium was added. On the third day, mesoderm lineage differentiation was initiated by removing 166 μL of old medium and applying 166 μL of RPMI 1640 medium containing insulin-free B-27 supplement, 4 μM CHIR99021, 1.25 ng/mL BMP4 and 1 ng/mL Activin A for 24 hours. The subsequent step involved removing 166 μL of medium followed by its replacement with fresh RPMI 1640 containing the insulin-free B-27 supplement. For specifying cardiac mesoderm, another medium change was performed, consisting of RPMI 1640 with insulin-free B-27 supplement and 3 μM Wnt-C59 for a final Wnt-C59 concentration of 2 μM, with an incubation period of 48 hours. Regular medium changes were continued every 48 hours, replacing 166 μL of medium with fresh RPMI 1640 and B-27 supplement.

### 7. MEA Field potential recording of cardiac organoids

The activity of cardiac organoids was recorded using a Maestro Edge system (Axion Biosystems) with 24-well CytoView MEA plates, each containing 16 electrodes. Cardiac organoids were prepared and transferred to individual wells. Organoid plating was done according to a previously reported method (26). Briefly, MEA plates were coated using Matrigel diluted in RPMI media (1:50) for at least one hour at 37°. After this time, the plates are washed and dried before transferring the organoids. Beating organoids were transferred one by one to precoated MEA plates with a small volume of media and maintained at 37°C and 5% CO₂ without disturbing for at least 4 to 5 hours. After this time, if needed, organoids were repositioned on top of the MEA chip to let them firmly attach for 8 to 24 hours in the incubator in 750 μl of incubation media. After the organoids are firmly attached, blebbistatin (10 μM) was added to the media and incubated at 37 °C for 2 to 4 hours before electrophysiological recordings.

Cell viability and spontaneous field potentials were acquired and recorded for 10 minutes using AxlS Navigator software version 3.10.1 (Axion Biosystems). Broadband signals were acquired at 12.5 kHz and high-passed filtered at 0.1 Hz for offline analysis. The beat detection threshold was set at 300 μV and a maximum period of 5 s. The beat period has been determined as the interval between two consecutive beatings on the same channel, and the field potential duration (FPD) was defined as the time between the start of the beating and the T wave, corresponding to the depolarization/repolarization (QT) interval. The FPD was corrected (cFPD) using the Bazett formula:

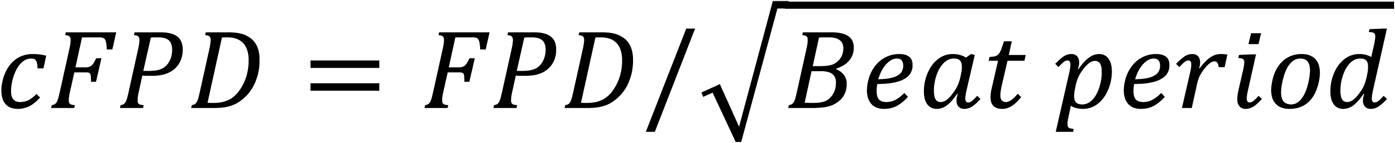

Data are presented as the mean ± SEM, and statistical analysis to determine differences between groups was performed using SigmaPlot 11.0 software (Grafiti LLC).

### 8. HLA Typing and Donor Selection

High-resolution HLA typing data were obtained from the SSCDR, which contains over 64,000 registered donors at the time of analysis. Population-level haplotype frequencies were estimated using the EM algorithm in Hapl-o-Mat v1.1 (as per Álvarez-Palomo et al., 2021) (27). Coverage was calculated iteratively by identifying the most frequent haplotypes, counting all matching individuals, and removing them from subsequent iterations. Matching criteria included HLA-A, - B, and -DRB1 loci, allowing matches on either of the two alleles per locus. Thirteen HLA-homozygous donors were ultimately selected to cover over 30% of the Saudi population.

### 9. Data Collection, Curation, and Management

Clinical and genetic information is securely stored at KAIMRC, accessible only to authorized SBiPSCs personnel. Each sample is assigned a unique, de-identified label, with the linking code maintained in a separate, restricted-access database. This approach ensures both patient privacy and regulatory compliance. Data sharing agreements with external collaborators require IRB approval and must conform to local and international data protection standards.

### 10. Cryopreservation and Biobanking Procedures

Each validated iPSC line is cryopreserved in 10% DMSO and mTeSR GMP-grade medium (STEMCELL Technologies). Vials are cooled using controlled-rate freezing protocols and stored in the vapor phase of liquid nitrogen. Duplicate vials are kept in separate cryostorage dewars as a failsafe measure. Vial labelling follows an internal coding system known only to SBiPSCs personnel. This labelling system allows linking each sample to its donor information, thus enabling traceability while preserving confidentiality.

## RESULTS

### SBiPSCs Patients-Derived Arm

KAIMRC–Western Region, which houses the SBiPSCs, is located within King Abdulaziz Medical City, a 751-bed tertiary-care complex of the Ministry of National Guard Health Affairs that serves the Western Region of Saudi Arabia. The campus integrates a broad spectrum of specialised facilities, including a Cardiology Centre, Neuroscience and Trauma Centre, a comprehensive Oncology Centre, a Bone-Marrow Transplantation Unit, and a network of primary-health-care clinics. In addition to a newly established a state-of-the-art tertiary children’s hospital that concentrates on complex paediatric conditions, many of which are genetic in origin. This clinical environment provides immediate access to diverse patient populations and advanced diagnostic resources, which provides an ideal setting for a national iPSC repository focused on disease modelling, drug discovery, and precision therapies.

To ensure high translational value of the bank, we implemented a clinic-guided enrolment strategy. Under this model, donors were recruited by physicians who have received briefing on the aims of the SBiPSCs and were instructed to nominate and prioritize patients with confirmed genetic diagnoses through targeted gene panels, whole-exome, or whole-genome sequencing. Most patients were recruited through the Neuroscience and Trauma Center, the Cardiology Center, and King Abdullah Specialist Children’s Hospital. When available, clinically unaffected relatives (e.g., parents or siblings) were co-recruited to generate genetically matched control lines.

To the day of writing this manuscript, the SBiPSCs has successfully established 37 characterized iPSC lines from 19 donors (Table 3). The composition of the bank is stratified into primary categories based on clinical pathology (Fig. 2). Neurological disorders constitute the predominant disease category within the SBiPSCs, collectively representing 48.6% of the repository (N=18 lines). This cohort is further stratified into Neurometabolic (16.2%; N = 6), Neurodegenerative (16.2%; N = 6), Neurodevelopmental (10.8%; N=4), and Neuromuscular (5.4%; N=2) disorders.

**Figure 2.**
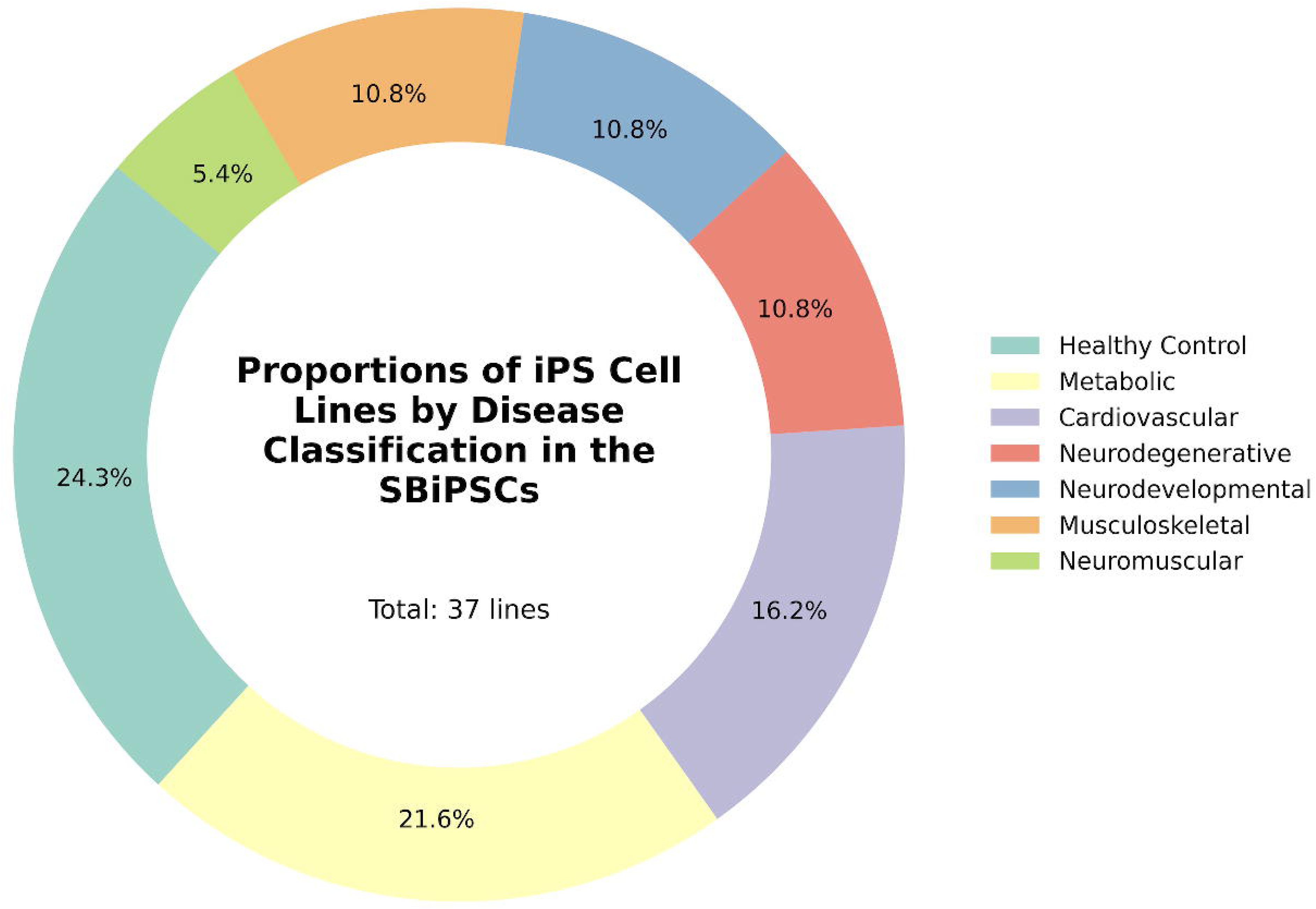
Distribution of iPSC lines by disease category in the Saudi Bank of Induced Pluripotent Stem Cells (SBiPSCs). Donut chart illustrating the proportional representation of the 37 fully characterized iPSC lines currently established in the SBiPSCs, stratified by primary clinical pathology. The repository includes healthy control lines (24.3%) and disease-specific lines encompassing Neurometabolic (16.2%), cardiovascular (16.2%), neurodegenerative (16.2%), neurodevelopmental (10.8%), musculoskeletal (10.8%), and neuromuscular (5.4%) disorders. Percentages indicate the fraction of total lines per category.

**Table 3.**
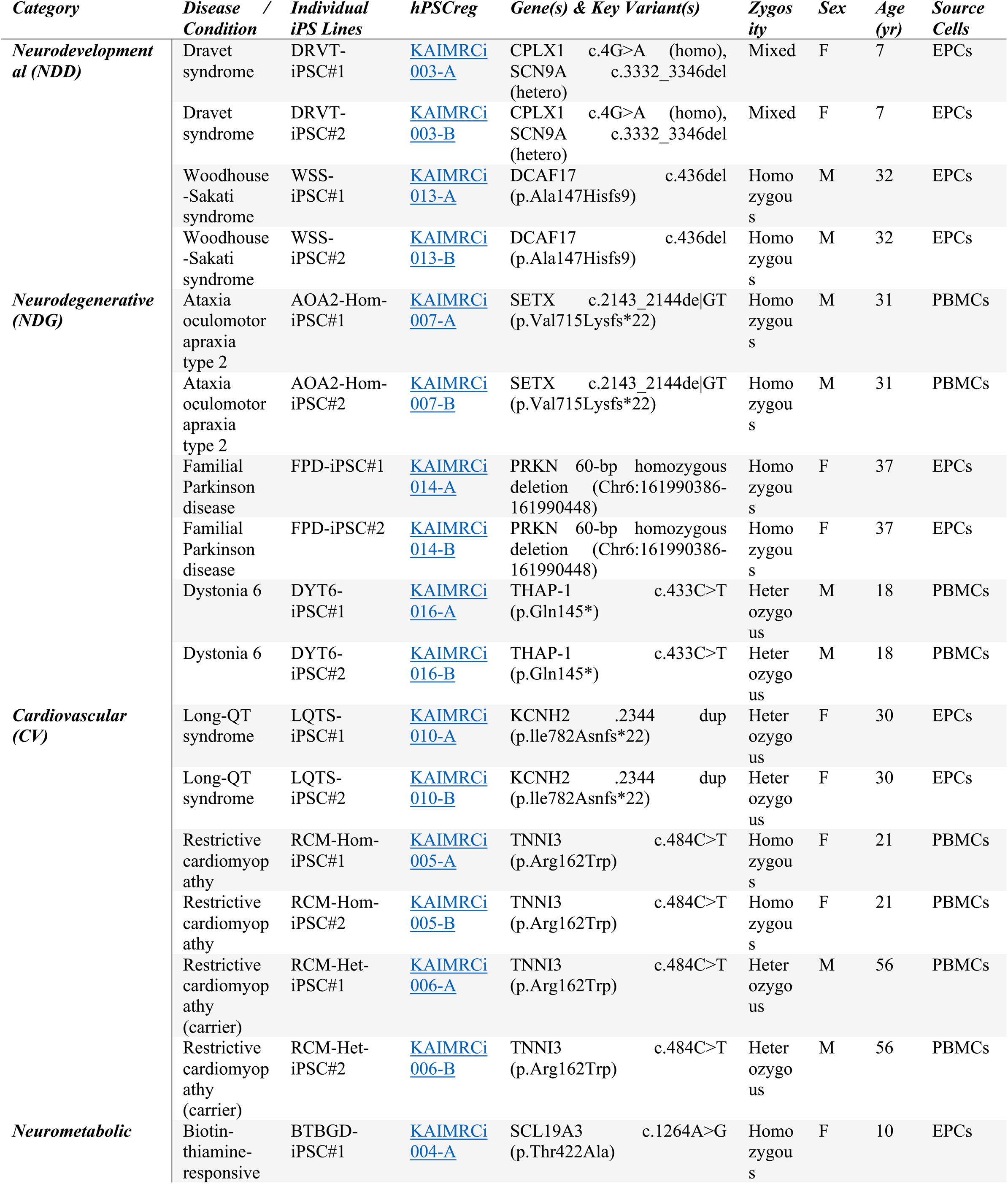
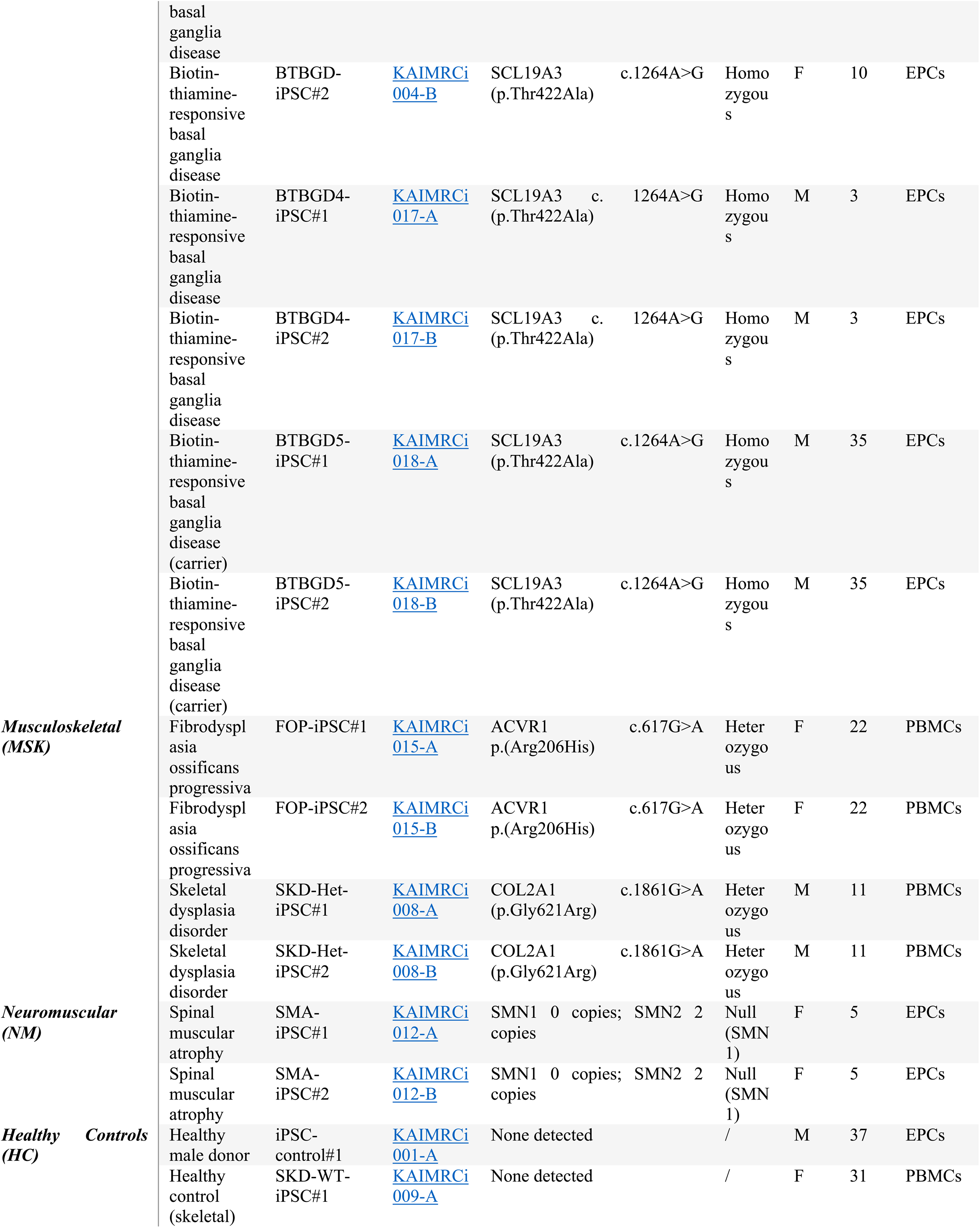
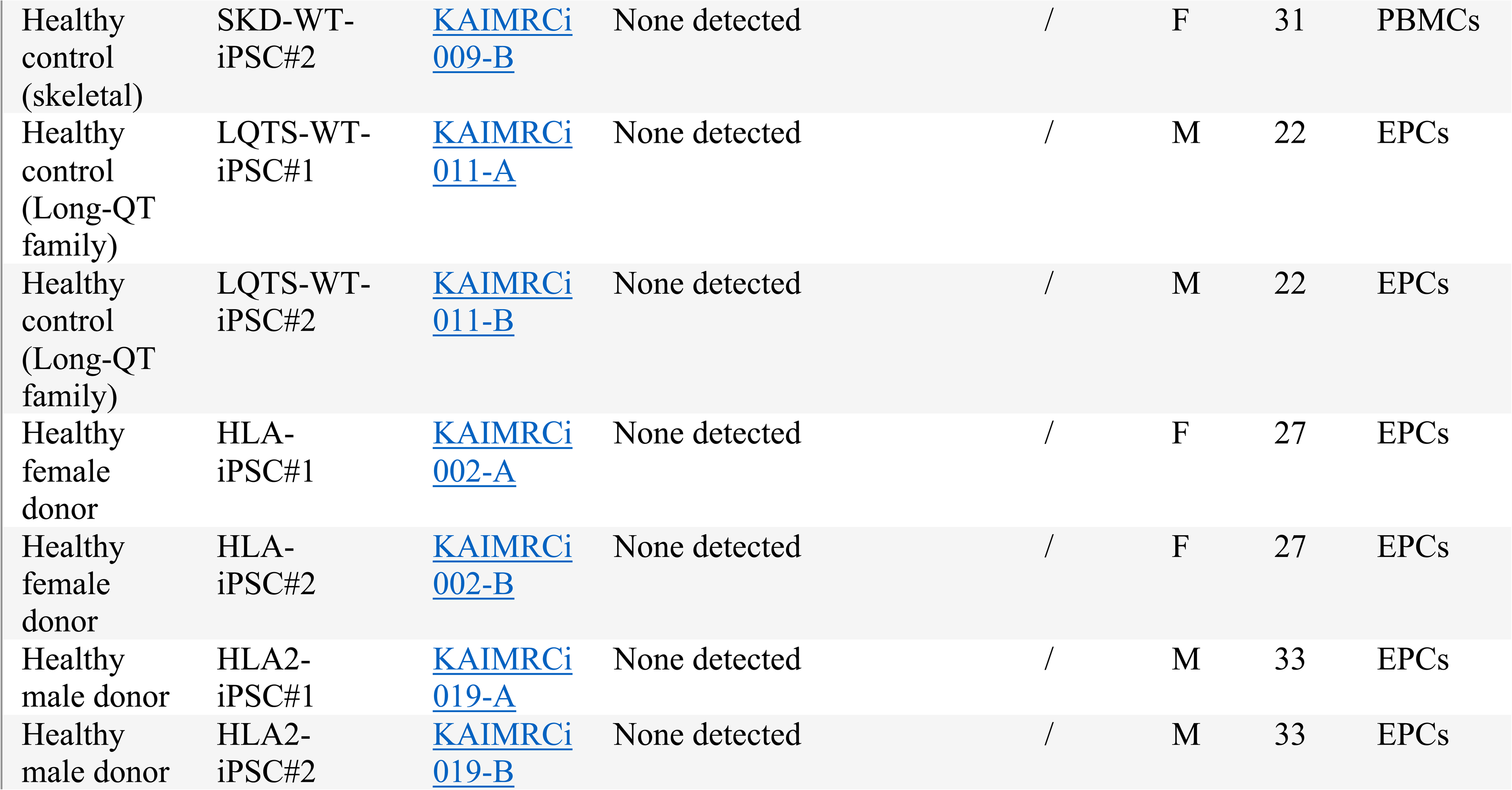
Distribution of characterized iPSC lines by disease category in the SBiPSCs.

Biotin-Thiamine-Responsive Basal Ganglia Disease (BTBGD) (*SLC19A3* c.1264A>G; p.Thr422Ala) (24) and Woodhouse-Sakati Syndrome (*DCAF17* c.436del; p.Ala147Hisfs*9).* To our knowledge, these represent the first reported iPSC lines generated for these specific disorders globally, providing unprecedented opportunity and resource for investigating their underlying molecular pathophysiology. Other key neurological lines include familial Parkinson’s disease (*PRKN* 60-bp homozygous deletion), Ataxia Oculomotor Apraxia type 2 *(SETX* c.2143_2144delGT; p.Val715Lysfs22), Dravet Syndrome (*CPLX1* c.4G>A / *SCN9A* c.3332_3346del) (23), Dystonia 6 (*THAP1* c.433C>T; p.Gln145*), and Spinal Muscular Atrophy (SMA) (*SMN1* 0 copies; *SMN2* 2 copies).

Cardiovascular disorders account for 16.2% of the collection (N=6 lines), encompassing models for channelopathies such as Long QT Syndrome (*KCNH2* c.2344dup; p.Ile782Asnfs*22) and structural defects like Restrictive Cardiomyopathy (*TNNI3* c.484C>T; p.Arg162Trp). Healthy controls currently constitute 24.3% of the bank, including lines from both unrelated healthy donors and unaffected familial controls (e.g., healthy siblings or parents). The inclusion of these familial controls is critical for generating isogenic-like comparisons that mitigate the confounding effects of genetic background in disease modelling studies.

Finally, Musculoskeletal disorders represent 10.8% (N=4 lines) of the bank, including lines for Fibrodysplasia Ossificans Progressiva (FOP) (*ACVR1* c.617G>A; p.Arg206His) and Skeletal Dysplasia (*COL2A1* c.1861G>A; p.Gly621Arg). This diverse and carefully curated collection serves as a robust platform for mechanistic studies and drug discovery, particularly for rare genetic diseases that have historically been underrepresented in global stem cell resources.

For each banked cell line, the SBiPSCs stores the patient’s molecular diagnostic report and the primary clinical diagnosis. Additional clinical variables may be accessed if needed through the treating physician, or through the BestCare™ platform. This access is restricted to authorized SBiPSCs personnel to protect the patients’ privacy. All donor identifiers are systematically replaced with coded labels that follow the standardized guidelines established by the Human Pluripotent Stem Cell Registry (hPSCreg). Internally, a laboratory information management system (LIMS) stratifies iPSC lines by disease category, gene mutation, and, when applicable, HLA haplotype.

### SBiPSCs HLA-Based iPSC Haplobank

Building upon our recent feasibility study (20), which established the computational framework for identifying optimal HLA-homozygous donors that maximize coverage within the Saudi Stem Cell Donor Registry (SSCDR), we have expanded our haplolines by adding an additional donor with the haplotype HLA-A*02:01 ∼ HLA-B*08:01 ∼ HLA-DRB1*03:01 that cover 2.4% of the Saudi population. As previously described (20), potential donors were stratified based on homozygosity for high-frequency HLA-A, -B, and -DRB1 haplotypes to maximize immunological compatibility across the Saudi population.

With this addition, the haplobank has expanded to include two fully validated homozygous donors, HLA1-iPSC lines derived in Alowaysi et al., 2023 (20) and HLA2, from which the iPSC lines described in this study were generated (Additional file 1: Fig. S1A-I). In total, four clonal iPSC lines have been generated (two clones per donor) that collectively cover 9% of the Saudi population. These lines were derived from erythroid progenitor cells to ensure genomic stability and have undergone rigorous characterization. The two recruited haplotypes and alleles are highly prevalent in the region. While the current lines are designated as research-grade and serve to support the development of allogeneic cell therapy workflows, KAIMRC is currently finalizing the commissioning of a dedicated Good Manufacturing Practice (GMP) facility. This facility will host the future expansion of this arm, enabling the derivation of clinical-grade iPSC lines compliant with regulatory standards for therapeutic applications.

### Representative Demonstration of the SBiPSCs Characterization Pipeline: Modeling Long QT Syndrome

To encourage donor participation, we optimized our reprogramming protocol for the SBiPSCs to use peripheral blood (PB) samples as the starting material. Unlike skin fibroblast-based methods, which require an invasive biopsy that may involve local anesthesia or suturing, PB samples collection is less intrusive and can be performed alongside routine clinical blood work.

We optimized our protocol so that PBMCs are electroporated on the same day of blood collection using non-integrating episomal plasmid reprogramming approach, which removes the need for intermediate expansion. Visible colonies typically emerge by day 14 and are ready for manual picking around day 21 post-transfection. For each donor, four to five colonies exhibiting typical iPSC morphology are selected, and two clones are expanded to passage 8 for complete quality control testing. If either clone exhibits abnormal karyotype or fails to express core pluripotency markers, a third backup clone is validated to ensure that each patient/donor ultimately contributes two lines that pass the characterization criteria. This rapid and standardized workflow has been successfully applied across the SBiPSCs patient-derived arm.

To demonstrate the SBiPSCs cell line characterization workflow and translational utility, in this report we present the case-study of a Saudi 30-year-old female patient with familial long QT syndrome. This patient was presenting with abnormal electrocardiography (ECG) with a prolonged QT interval of 476 milliseconds. She was referred to the SBiPSCs by her treating physician, and after consenting, her blood sample was collected with her 22-year-old unaffected brother to serve as a control. Diagnostic Whole Exome Sequencing (WES), that had been previously requested by the treating physician (performed by the clinical Molecular Diagnostics laboratory), identified a heterozygous mutation in the *KCNH2* gene (c.2344dup p.(Ile782Asnfs*22)). The mutation is a known pathogenic variant associated with LQTS Type 2, as previously reported by Bhuiyan et al. in 2009 (28). This mutation introduces a premature stop codon, predicted to result in nonsense-mediated decay or a truncated non-functional protein. Here, we confirmed this finding through Sanger sequencing of the patient’s peripheral blood cells and derived iPSCs, as shown in Fig. 3D.

**Figure 3.**
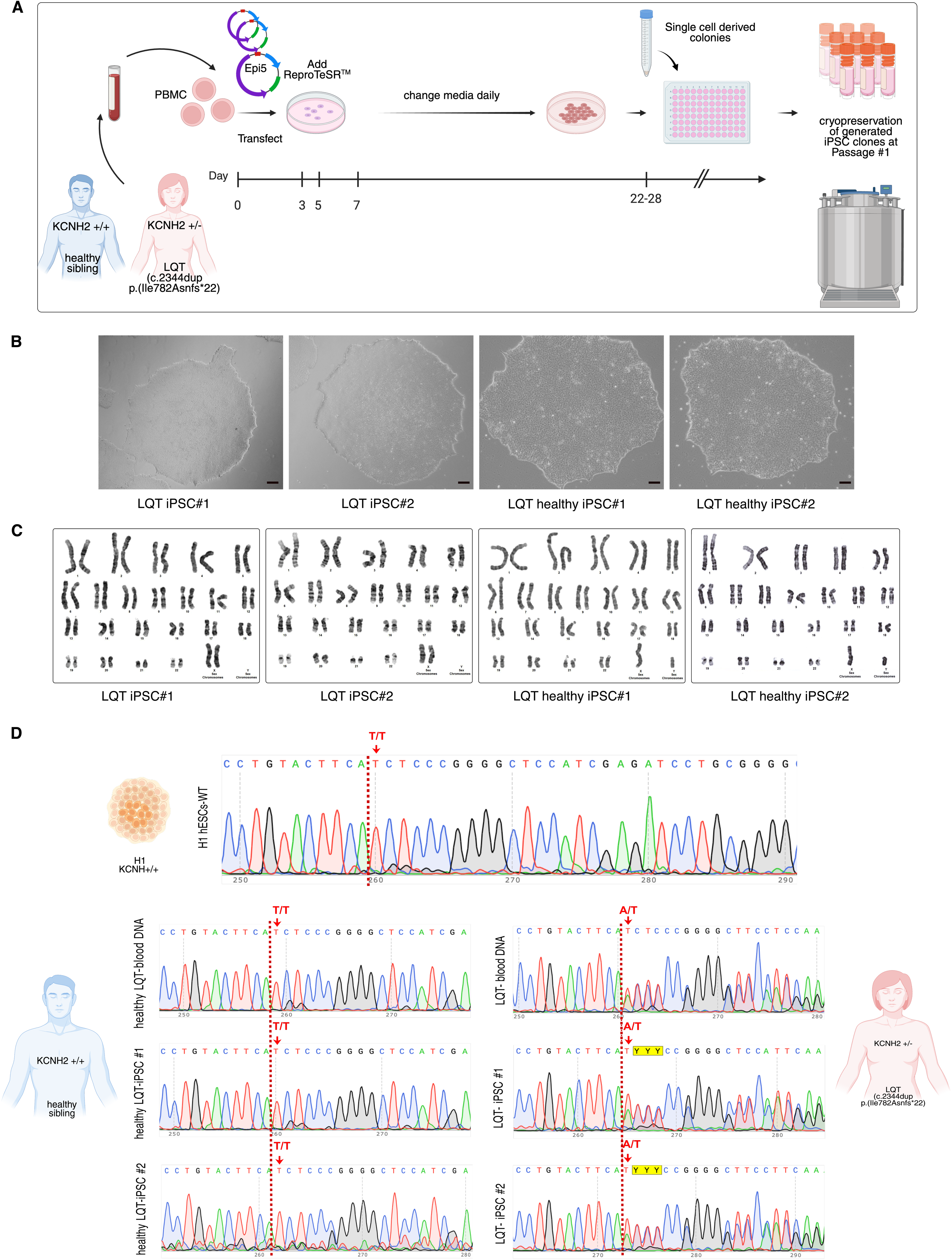
The reprogramming and generation of LQTS-iPSCs. A) A Schematic illustrating the sample collection and outlining iPSCs reprogramming methods. B) Representative images of LQTS-iPSC clones, which are shown in tightly packed groups with distinct borders. Scale bar, 100 μm. C) Standard G-banded karyotyping confirms that LQTS-iPSCs #1 and 2 maintain normal chromosomal content, specifically 46, XX and normal chromosomal profile for LQTS healthy-iPSCs #1 and 2, specifically 46, XY. D) The electropherogram record from Sanger sequencing shows KCNH2 mutations in the patient’s peripheral blood cells and the LQTS-iPSC #1 and 2 lines.

Four clonal lines were established in total, two from the LQTS patient and two from the healthy control. Approximately 20 days after transfection of the EPCs with the episomal plasmids, several colonies emerged exhibiting characteristic features of embryonic stem cells, including well-defined edges, dense cell packing, prominent nucleoli and high nucleus to cytoplasm ratio (Fig. 3A, B). These colonies were manually picked and expanded and cryopreserved at the KAIMRC facility. Based on the morphology and rate of cell division, two clones per donor were selected for further characterization.

Chromosomal analysis revealed that both patient lines (LQTS-iPSCs #1 and #2) maintained a normal female karyotype (46, XX), while the control lines (LQTS healthy-iPSCs #1 and #2) displayed a normal male karyotype (46, XY) (Fig. 3C). Short tandem repeat (STR) analysis was performed to confirm the donor identity, which confirmed that the iPSC lines were genetically identical to the respective donor’s PBMCs (Additional file 2: Fig. S2B). Furthermore, routine screening confirmed that all generated iPSC lines were free of mycoplasma contamination (Additional file 2: Fig. S2C). Additionally, molecular analysis of genomic DNA and RNA extracts confirmed the absence of blood-borne pathogens, including Human Immunodeficiency Virus (HIV), Hepatitis B Virus (HBV), and Hepatitis C Virus (HCV). These stringent tests are quality control measures implemented across the SBiPSCs repository to ensure that all banked iPS cell lines are karyotypically normal (unless the patient presents with an aneuploidy) and free of infectious agents.

The clones were expanded and tested for episomal vector persistence from passage six onward, confirming complete loss of reprogramming plasmids between passages twelve and sixteen (Additional file 2: Fig. S2A). Pluripotency was assessed by evaluating OCT4, NANOG, and SOX2 expression using flow cytometry, immunofluorescence, and RT-qPCR. Flow cytometry analysis showed that more than 95% of cells expressed OCT4 and NANOG, while 97% were positive for SOX2 (Fig. 4A). Immunofluorescence analysis confirmed strong expression of pluripotency markers in the iPSC lines (Fig. 4B). RT-qPCR further revealed a marked upregulation of *POU5F1*, *NANOG*, and *SOX2* transcripts compared with the H1 hESC positive control (Fig. 4C). The competency of the LQTS-iPSC lines to differentiate into mesoderm, endoderm, and ectoderm was tested via direct in vitro differentiation. We detected an upregulation of lineage-specific markers and a downregulation of the pluripotency markers *OCT4* and *NANOG* across all lineages by RT-qPCR. Ectoderm differentiation was confirmed by the expression of the central nervous system neural progenitor markers *PAX6* and *NESTIN*. Mesoderm differentiation was evidenced by increased expression of a member of the T-box family of transcription factors (*Brachyury*) and a caudal-type homeobox protein 2 (*CDX2*). The upregulation of endodermal markers, including the zinc-finger transcription factor (*GATA4*) and the SRY-Box transcription factor 17 (*SOX17*), was confirmed in LQTS-iPSC lines and H1 hESC control (Fig. 5A). Following in vitro differentiation, the cells were analyzed by immunocytochemistry for germ layer markers. Positive staining was observed for PAX6, Desmin, and SOX17, indicating successful differentiation into ectodermal, mesodermal, and endodermal lineages, respectively (Fig. 5B). Upon confirming the pluripotent status of the generated lines, the LQTS-iPSCs were officially registered in the Human Pluripotent Stem Cell Registry: https://hpscreg.eu/cell-line/KAIMRCi010-B, https://hpscreg.eu/cell-line/KAIMRCi011-A and https://hpscreg.eu/cell-line/KAIMRCi011-B

**Figure 4.**
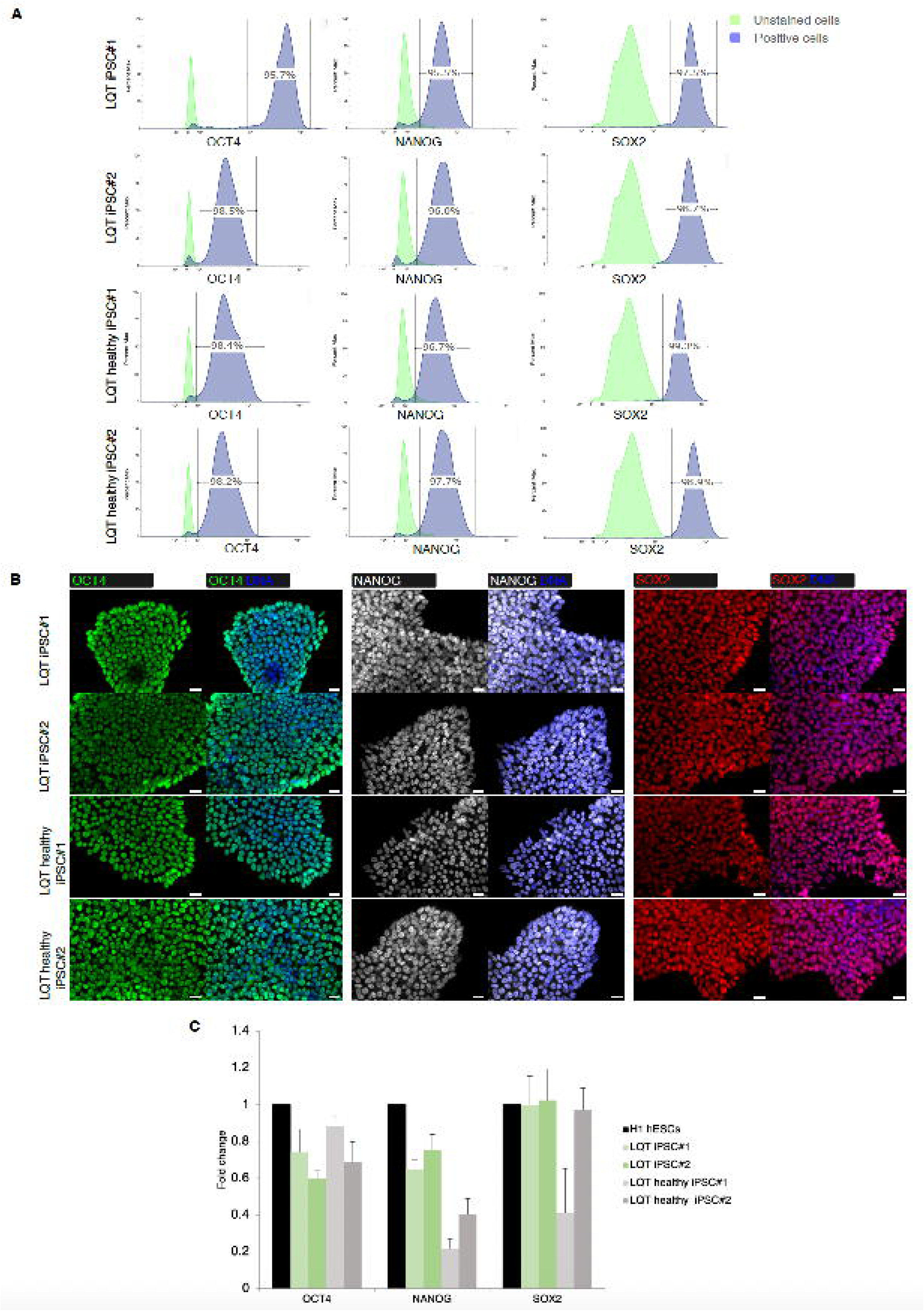
Confirmation of pluripotency markers in the generated LQTS-iPSC lines. A) Flow cytometry histograms illustrate the expression of pluripotency markers OCT4, NANOG, and SOX2 in LQTS-iPSCs. B) Immunofluorescence staining shows the presence of pluripotency markers OCT4 in green, NANOG in grey, and SOX2 in red, with nuclei counterstained with DAPI (blue). Scale bars, 100 μm. C) Real-time PCR (q-PCR) showing the mRNA expression levels of pluripotency markers for the LQTS-iPSC lines illustrated as a fold change relative to H1 hESC. The data is represented with bars indicating the median ± standard deviation.

### Efficient generation and MEA-based functional characterization of cardiac organoids from LQT-iPSCs

To demonstrate the translational utility of the SBiPSCs repository for disease modeling, we assessed the ability of the derived Long QT Syndrome iPS cell lines to differentiate into three-dimensional (3D) cardiac organoids. Given that cardiac channelopathies such as Long QT Syndrome manifest primarily through electrical instability in the myocardium, the generation of functionally active cardiac tissue allows for the studying of disease mechanisms *in vitro*.

Using a defined stepwise differentiation protocol (25), both patient (LQTS iPSC#1) and control (LQTS Healthy iPSCs#1) lines were successfully differentiated to form cardiac organoids (Fig. 6A). Differentiation efficiency was robust across all lines, yielding 3D aggregates that exhibited spontaneous contractions visible under bright-field microscopy by day 10 (Fig. 6B) (Additional file 3: # movie 1) (Additional file 4: # movie 2). Immunofluorescence analysis revealed widespread and organized expression of the cardiomyocyte-specific structural protein Cardiac Troponin T (TNNT2) (Fig. 6C). This demonstrates that the iPS cell lines banked within the SBiPSCs possess the capacity to generate complex, functional tissues suitable for disease modeling.

**Figure 5.**
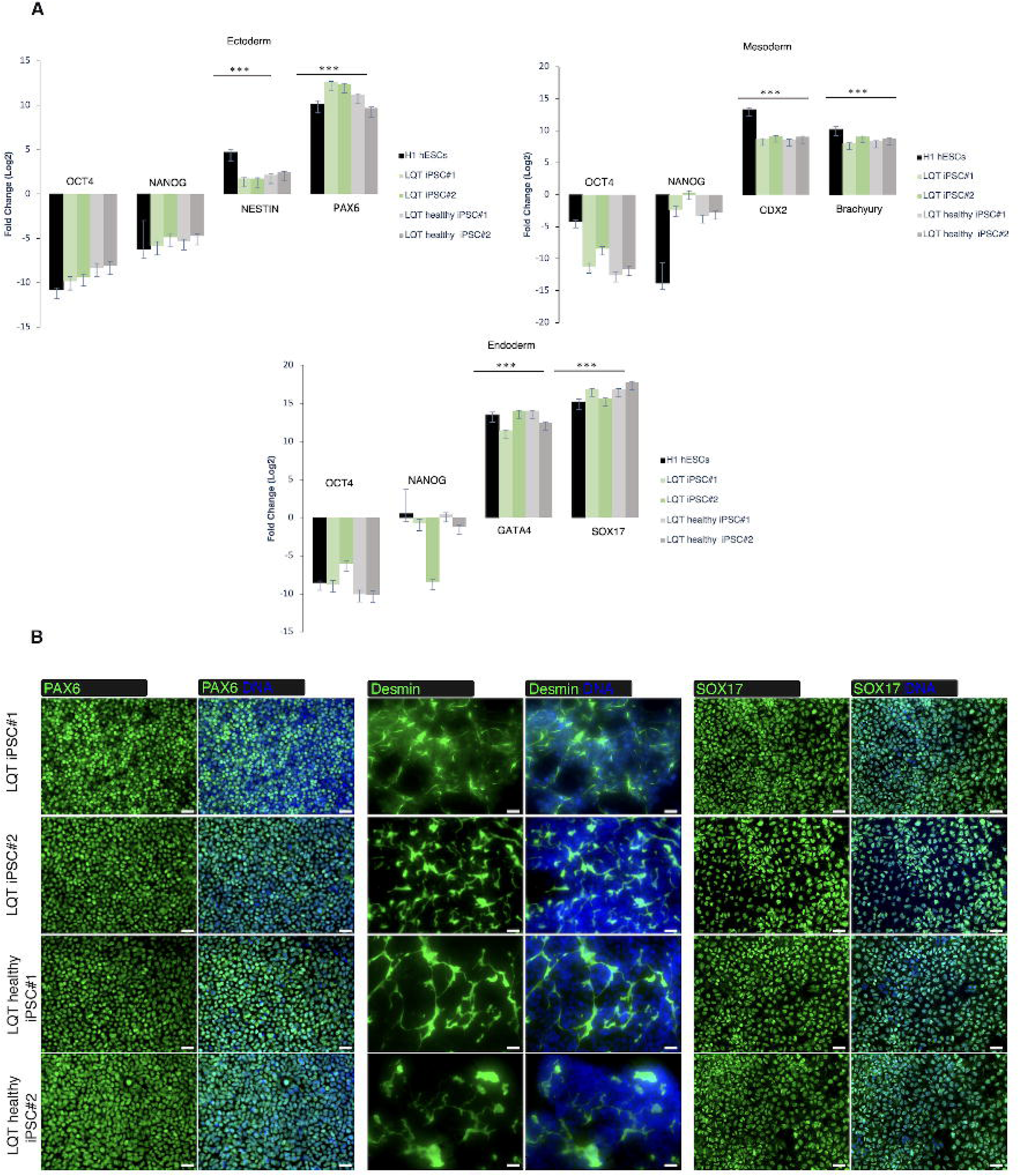
Assessing the potential of the generated LQTS-iPSC lines to differentiate into the three germ layers. A) qPCR data showing the mRNA expression levels of lineage-specific markers consisting of ectoderm (NESTIN and PAX6), mesoderm (CDX2 and Brachyury), and endoderm (GATA4 and SOX17). Data are shown as fold changes in comparison to undifferentiated cells. The data bars indicate the median ± standard deviation, with statistical significance denoted by Student’s t-tests, *p<0.05. B) Immunofluorescence staining displays the expression of PAX6 (ectoderm), Desmin (mesoderm), and SOX17 (endoderm) in tissues derived from the three embryonic germ layers (Green) in generated LQTS-iPSCs, with nuclei counterstained using DAPI (blue). Scale bar, 100 μm.

**Figure 6.**
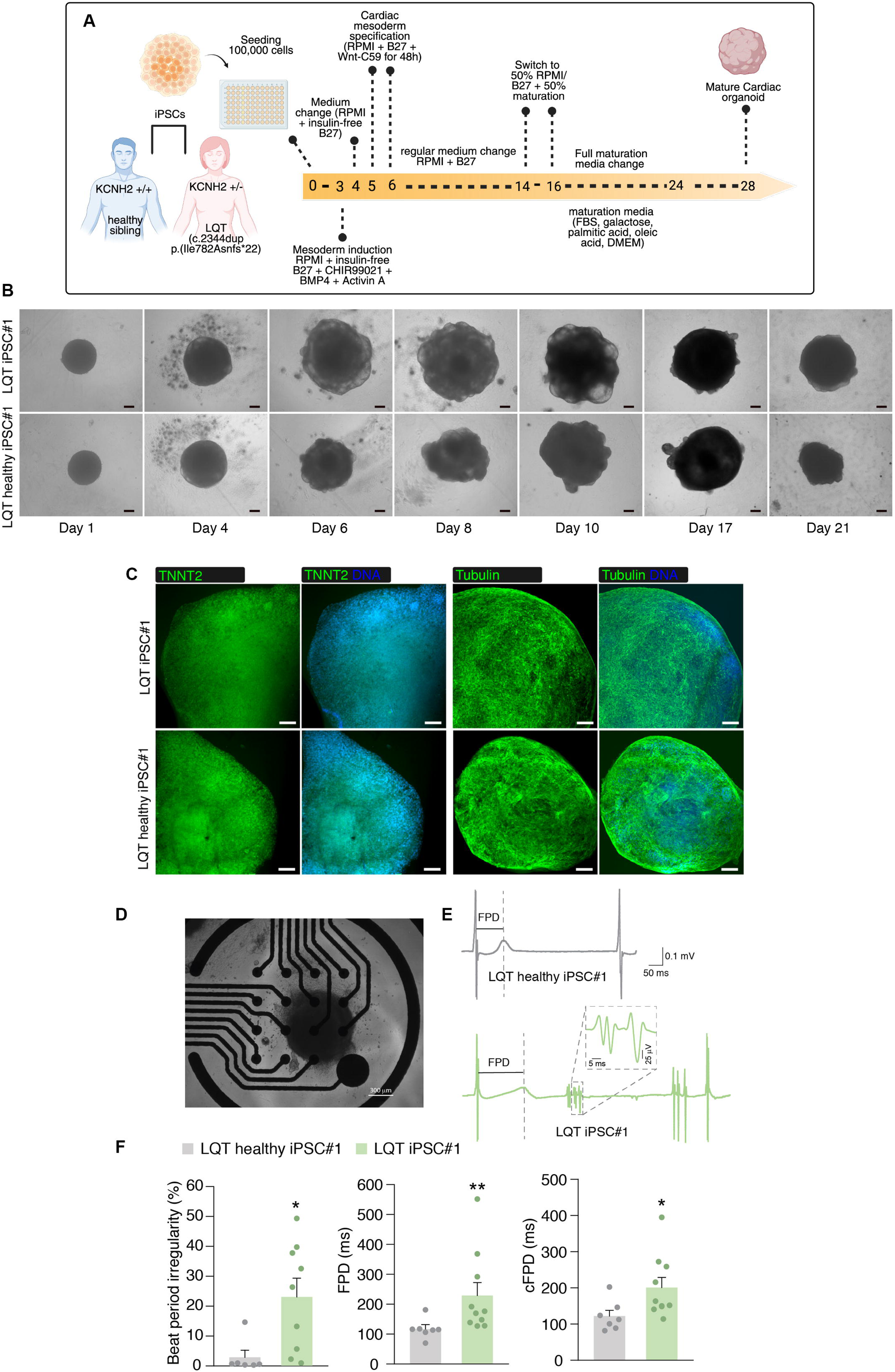
Differentiation of LQTS-iPSCs into heart organoids. A) A diagram showing the methods and timeline for differentiating LQTS-iPSCs into cardiac organoids. B) Bright-field images of cardiac organoids from day 1 to day 18. Scale bar, 100 μm. C) Immunofluorescence staining shows the presence of cardiomyocyte marker TNNT2 (Green) in the created LQTS-iPSC-derived cardiomyocyte, with the nuclei stained using DAPI (blue). Scale bar, 100 μm. D). Representative image of an iPSC-derived cardiac organoid placed on a MEA chip. Scale bar 300 μM. E) Representative field potential traces recorded from LQTS healthy iPSC#1-derived cardiac organoids (top, gray) and Long QT Syndrome (LQTS) iPSC#1-derived cardiac organoids (bottom, green). The Field Potential Duration (FPD) is indicated by dashed lines in both traces. The LQTS trace shows characteristic prolongation of the field potential duration and an irregular repolarization phase. The inset provides an expanded view of the LQTS field potential signal showing complex signals interpreted as out-of-rhythm afterdepolarizations. F) Quantitative analysis of MEA recordings comparing LQTS healthy cardiac organoids (gray bars) and LQTS organoids (green bars). The beat period irregularity (left), field potential duration (FPD) (center), and the corrected field potential duration (cFPD) (right) are significantly increased in LQTS cardiac organoids. *p < 0.05; **p < 0.01. Mann-Whitney U. Data are presented as mean ± SEM of 7 to 8 individual organoids.

To further validate the utility of the iPS-derived cardiac organoids for disease modeling, we performed microelectrode array (MEA) analysis to compare the electrophysiological profiles of organoids derived from the LQTS patient versus the healthy sibling to assess the model’s fidelity. Spontaneous field potentials were recorded with broadband signals acquired at 12.5 kHz and high-pass filtered at 0.1 Hz to isolate relevant field potential waveforms (Fig. 6D). The analysis showed that cardiac organoids derived from LQTS iPSC#1 exhibited significantly prolonged Field Potential Durations (FPD) compared to those derived from the LQTS Healthy iPSC#1 (Fig. 6E). The healthy sibling’s organoids displayed normal repolarization kinetics, while the patient-derived organoids showed a marked extension in the interval between depolarization and the T-wave terminus.

When corrected for beat rate using the Bazett formula, the patient-derived organoids displayed a significantly increased cFPD (*p* < 0.05), which reflects the prolonged QTc interval observed in the donor’s clinical ECG. Furthermore, quantitative analysis of beat period irregularity demonstrated a higher incidence of arrhythmic events in the LQTS iPSC#1 organoids compared to controls (Fig. 6F). This ability to faithfully recapitulate the donor’s specific channelopathy phenotype, both in terms of repolarization delay and arrhythmic susceptibility, confirms the fidelity of the cardiac organoids model. This functional validation study highlights the potential of the SBiPSCs platform as a robust resource for mechanistic studies, personalized drug screening, and precision medicine applications for the population in the region.

## DISCUSSION

The realization of precision medicine relies heavily on the availability of diverse genomic resources that reflect the full spectrum of human mutation variability (1,2). However, the existing global iPSC resources and infrastructures are characterized by a stark “genomic divide,” with the vast majority of available cell lines being derived from European, East Asian, and North American populations (3–5). This imbalance limits the applicability of current disease models and drug screening platforms to underrepresented groups, particularly those in the MENA region (2,4,5). The establishment of the Saudi Bank of Induced Pluripotent Stem Cells aims to address this critical gap. To our knowledge, SBiPSCs represents the first centralized national iPSC repository in the Middle East, that provide the global scientific community with access to a distinct genetic landscape that is affected by high rates of consanguinity and large family structures, making it well-suited for dissecting the molecular mechanisms of rare and recessive disorders (15,16).

A defining characteristic of the SBiPSCs is its integration within a tertiary care clinical framework, which facilitates the targeted recruitment of patients with clearly defined, often rare, genetic etiologies. Major biobanking initiatives, such as the European Bank for induced Pluripotent Stem Cells (EBiSC), have reported that patient recruitment and depositor participation remain significant operational challenges (10). In contrast, our clinic-guided strategy, embedded within the King Abdulaziz Medical City, streamlines donor identification and consent, and it ensures a steady recruitment of potentially informative genotypes.

This approach is exemplified by our establishment of iPSC lines for Woodhouse-Sakati Syndrome (*DCAF17*) and Biotin-Thiamine-Responsive Basal Ganglia Disease (*SLC19A3*) (24). Because consanguineous populations often harbor homozygous loss-of-function mutations in a natural “human knockout” context, these lines offer a valuable resource for functional genomics, allowing researchers to interrogate gene function in a relevant human background without the potential off-target effects associated with user-induced gene editing (24,30).

To demonstrate the translational utility of our repository, we utilized Long QT Syndrome (LQTS) as a benchmark for our characterization pipeline. Cardiac channelopathies are challenging to model in heterologous systems due to the lack of the complex electromechanical architecture found in native cardiomyocytes (17,19). Our results indicate that SBiPSCs-derived lines can be efficiently differentiated into 3D cardiac organoids that recapitulate the donor’s specific electrophysiological phenotype. The observation of a significantly prolonged field potential duration (FPD) in patient-derived organoids, alongside the generation of a control cardiac organoids from a healthy sibling, supports the fidelity and utility of our banked iPS cell lines. It is important to note that the microelectrode array analysis presented here was conducted as a single experimental set to demonstrate the translational utility of the lines for disease modeling, rather than as a comprehensive mechanistic investigation of the channelopathy itself. Nevertheless, the demonstrated recapitulation of the disease phenotype seen in clinical ECG findings *in vitro* suggests that the repository supports the generation of physiologically relevant tissues suitable for drug toxicity screening.

In parallel with disease modeling, the SBiPSCs is establishing a foundation for future regenerative therapies through its HLA-homozygous haplobank. Our strategy aligns with initiatives like Japan’s CiRA project, which demonstrated that a small number of carefully selected homozygous donors can provide immunological matching for a significant segment of the population (31). By examining the high resolution HLA database of the Saudi Stem Cell Donor Registry, we have achieved ∼9% population coverage with two donors HLA1-iPSC lines (20) and HLA2, from which the iPSC lines described in this study were generated. It is worth noting that these haplolines are research-grade and are currently being utilized in multiple projects to develop off-the-shelf iPSC-based therapies. KAIMRC is currently in the final stages of launching its state-of-the-art GMP-grade cell processing center in Riyadh, which will host the expansion of HLA-based banking arm of the SBiPSCs.

Despite these advances, several limitations should be acknowledged. The current repository comprises 37 iPSC lines, which remains modest when compared with large, established international consortia. However, this banking initiative was formally launched in 2023 and represents an early-phase national infrastructure effort rather than a mature collection. Importantly, the repository is embedded within a strategically funded national framework aligned with Saudi Vision 2030 and the National Biotechnology Strategy, which prioritize the development of population-representative biomedical resources, precision medicine, and cell-based therapeutics. Sustained institutional and governmental investment is in place to support systematic donor recruitment, large-scale derivation, and long-term expansion of the bank.

Another potential limitation is that functional validation in the present study focused on cardiac organoid modeling. This choice was deliberate, as Long QT Syndrome provides a robust, quantifiable electrophysiological phenotype that is well suited for benchmarking disease modeling fidelity. Nevertheless, cardiomyocytes represent only one of many clinically relevant lineages. Ongoing and planned efforts are directed toward extending validation across additional germ-layer derivatives and disease contexts, including neural, metabolic, and hematopoietic lineages, using standardized differentiation pipelines and lineage-specific functional assays. As the repository expands, these multi-lineage validation workflows will be applied systematically to ensure broad utility of the bank for diverse disease-modeling and translational applications.

In summary, the SBiPSCs provides a vital addition to the global stem cell landscape. By combining a distinct demographic niche with rigorous quality control, the bank is positioned to support the discovery of population-specific disease mechanisms and facilitate the development of inclusive, precision therapeutics. As the repository expands, it will serve as a resource for regional biotechnology and a model for establishing similar infrastructures in other underrepresented populations.

## Conclusions

The SBiPSCs represents the first national iPSC repository in the Middle East and North Africa, directly addressing the longstanding global underrepresentation of Middle Eastern populations in stem cell resources. Through the successful generation of patient-specific iPSC lines, SBiPSCs establishes a reliable and reproducible workflow for disease modeling and functional validation. The initiative will further lay the groundwork for HLA-guided iPSC banking under Good Manufacturing Practice (GMP) standards, enabling future applications in regenerative and cell-based therapies. By integrating patient recruitment, biobanking, organoid modeling, and genome editing within a unified platform, SBiPSCs strengthens Saudi Arabia’s capacity for precision medicine and positions the Kingdom as a regional leader in translational stem cell research. By combining a distinct demographic focus with rigorous scientific validation and a clear strategic vision, SBiPSCs constitutes a vital addition to the global stem cell landscape, serving as a cornerstone for inclusive precision therapeutics and a scalable model for advancing biomedical research in underrepresented populations worldwide.

## Abbreviations

iPSC(s): induced pluripotent stem cell(s)
SBiPSCs: Saudi Bank of Induced Pluripotent Stem Cells
KAIMRC: King Abdullah International Medical Research Center
MENA: Middle East and North Africa
LQTS: Long QT Syndrome
PBMC(s): peripheral blood mononuclear cell(s)
HLA: human leukocyte antigen
GMP: Good Manufacturing Practice
ECG: electrocardiogram
hESC(s): human embryonic stem cell(s)
QC: quality control
CiRA: Center for iPS Cell Research and Application
EBiSC: European Bank for induced pluripotent Stem Cells
JCRB: Japanese Cancer Research Resources Bank
SSCDR: Saudi Stem Cell Donor Registry
SD: standard deviation
RT-qPCR: reverse transcription quantitative polymerase chain reaction.

## Acknowledgements

We thank King Abdullah International Medical Research Center (KAIMRC) and the Saudi Stem Cell Donor Registry (SSCDR) for facilitating the initiation of HLA-based iPSC banking. We gratefully acknowledge the support of Pfizer through the Saudi Economic Participation Program, administered by the Local Content and Government Procurement Authority (LCGPA), for funding the patient-derived arm of the SBiPSCs project. This national program is designed to channel part of the financial value of government-related contracts with foreign entities into high-impact national initiatives. A core focus of the Economic Participation Program is to support applied research and development projects within the Kingdom that align with strategic national priorities and the Kingdom’s Vision 2030, such as localizing advanced technologies, enhancing knowledge transfer, and building local scientific capabilities. The establishment of the SBiPSCs exemplifies this mandate by creating a regional centralized iPSC platform for disease modelling, drug discovery, and future cell-based therapies tailored to the Saudi population. Finally, we extend our deepest appreciation to all current and future donors for their valuable contributions to this national biobanking initiative.

## Author contributions

K.A. conceived the study, directed the project, secured funding, and wrote the manuscript. M.H., M.A., and Y.F. performed and designed experiments, analyzed data, and participated in writing the manuscript. M.B., M.A.-S., G.R., S.Z., N.A., L.Aljahdali, and L.Aljuid performed experiments related to iPSC generation and validation and generated and analyzed data. G.H.-L. performed MEA experiments and analyzed data under the supervision of P.M. D.A. performed STR analysis. H.A. performed karyotyping. F.H. managed the genetic testing of patients and data curation. A.Alaskar and D.J. curated and provided the SSCDR database. S.M., A.H., and A.Attar contributed to patient selection and recruitment. J.T. and D.G.-C. provided resources and contributed to the writing and editing of the manuscript. All authors read and approved the final manuscript.

## Availability of data and materials

All the presented data are available for consultation.

## Declarations

### Ethics approval and consent to participate

This project titled “Establishment of the Saudi Bank of Induced Pluripotent Stem Cells (SBiPSCs) A National Platform for iPSC-Based Research and Therapy” was approved by the institutional review board (IRB) and research ethics committee at the Ministry of National Guard for Health Affairs on December 3rd, 2020 (study numbers RJ20/134/J& RJ22J/060/03). Approved informed consent was obtained on October 22, 2021. This study was conducted in accordance with the ethical standards laid down in the 1964 Declaration of Helsinki and its later amendments.

## Figure legends

**Additional file 1** Figure 1S HLA-guided derivation and characterization of homozygous iPSC lines for allogeneic banking in the Saudi population. (A) Schematic overview illustrating HLA-guided donor selection, reprogramming of peripheral blood mononuclear cells (PBMCs) into two HLA-homozygous iPSC lines, and the resulting ∼9% population coverage within the Saudi population, supporting the feasibility of HLA-based iPSC banking for future allogeneic applications. Conceptual schematic created with the assistance of AI-based visualization tools and finalized by the authors. (B) Representative phase-contrast images showing that HLA2-iPSCs display typical human embryonic stem cell (ESC)-like morphology. (C) Normal karyotype analysis demonstrating a male (46, XY) chromosomal profile. (D) Clearance of episomal reprogramming plasmids (EPi5) in derived iPSC lines by passage 9. (E) RT-qPCR analysis showing expression of pluripotency markers at levels comparable to H1 human embryonic stem cells (hESCs). Bars are median ± std of 3 biological replicates for each sample. (F) Immunocytochemistry (ICC) confirming expression of core pluripotency markers. Scale bar=200 μm (G) Flow cytometry (FACS) analysis demonstrating robust expression of pluripotency-associated surface markers. (H) Trilineage differentiation assessed by ICC, confirming ectoderm (PAX6), endoderm (SOX17), and mesoderm (Desmin) marker expression; nuclei are counterstained with DAPI (blue). Scale bar=200. (I) Short tandem repeat (STR) profiling demonstrating genetic identity between HLA2-PBMCs and the corresponding derived iPSC lines.

**Additional file 2** Figure 2S. A) Representative image of gel electrophoresis from PCR showing the absence of the five episomal plasmids in the LQTS-iPSC lines. B) Short Tandem Repeat (STR) profiling guaranteed the genetic identity between the established iPSC lines and the donor PBMCs. C) Image of mycoplasma test in LQTS-iPSC lines illustrating the absence of contamination.

**Additional file 3 Movie#1.** Spontaneous beating of LQTS healthy-cardiac organoids appeared as early as day 10 of the differentiation protocol. The video was recorded at 10X magnification.

**Additional file 4 Movie#2.** Spontaneous beating of LQTS -cardiac organoids appeared as early as day 10 of the differentiation protocol. The video was recorded at 10X magnification.

