## Supplementary Figure 1 for "Establishment of the Saudi Bank of Induced Pluripotent Stem Cells (SBiPSCs): A National Platform for iPSC-Based Research and Therapy"

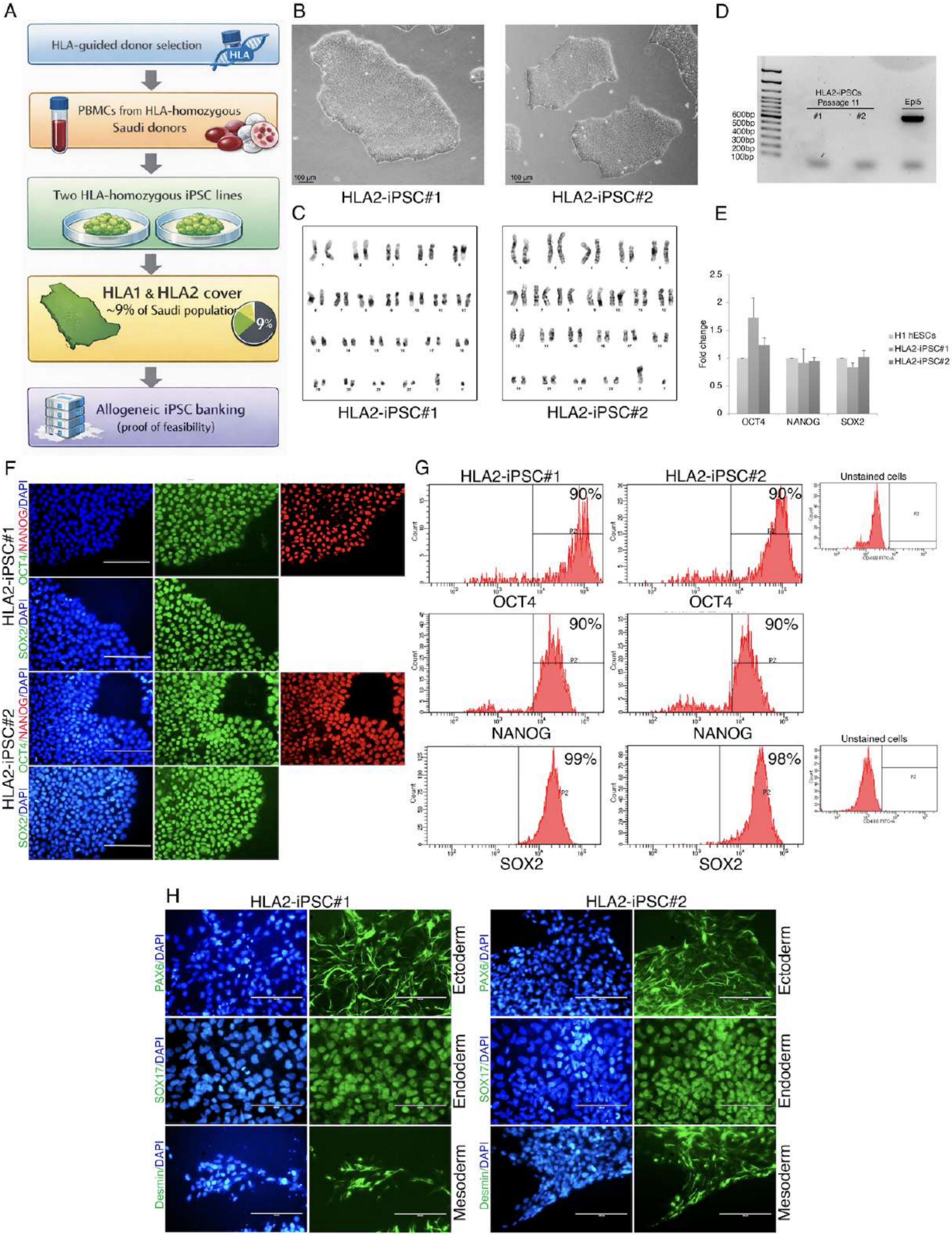


I

| STR markers | D3S1358 | vWA | D16S539 | AMEL | D8S1179 | D21S11 | D18S51 | D19S433 | TH01 | FGA | D5S818 | D13S317 | D7S820 | D10S1248 | D1S1656 | D2S1338 | CSF1PO | TPOX | Yindel | DYS391 | D2S441 |
|---|---|---|---|---|---|---|---|---|---|---|---|---|---|---|---|---|---|---|---|---|---|
| PBMCs | 16/17 | 18/18 | 9/9 | X/Y | 11/13 | 30/30 | 11/14 | 13.2/13.2 | 6/7 | 21/21 | 10/10 | 12/14 | 9/11 | 14/15 | 12/16 | 20/20 | 11/11 | 9/12 | 2/2 | 9/9 | 11/12 |
| HLA2-iPSC#1 | 16/17 | 18/18 | 9/9 | X/Y | 11/13 | 30/30 | 11/14 | 13.2/13.2 | 6/7 | 21/21 | 10/10 | 12/14 | 9/11 | 14/15 | 12/16 | 20/20 | 11/11 | 9/12 | 2/2 | 9/9 | 11/12 |
| HLA2-iPSC#2 | 16/17 | 18/18 | 9/9 | X/Y | 11/13 | 30/30 | 11/14 | 13.2/13.2 | 6/7 | 21/21 | 10/10 | 12/14 | 9/11 | 14/15 | 12/16 | 20/20 | 11/11 | 9/12 | 2/2 | 9/9 | 11/12 |
