## Supplementary Figure 2 for "Establishment of the Saudi Bank of Induced Pluripotent Stem Cells (SBiPSCs): A National Platform for iPSC-Based Research and Therapy"

**A**

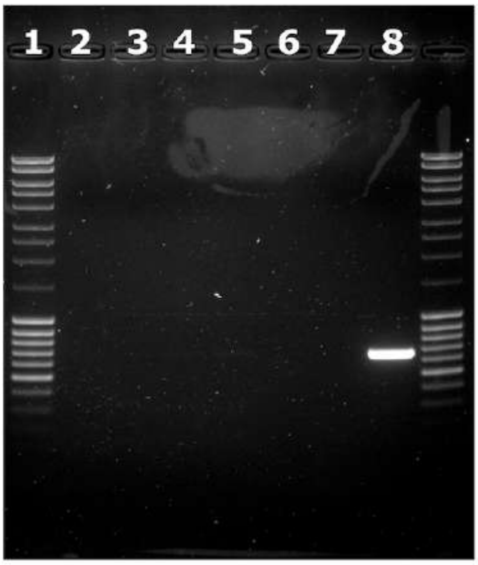


1-100pb ladder
2-LQT iPSC#1
3-LQT iPSC#2
4-LQT healthy iPSC#1
5-LQT healthy iPSC#2
6-Space
7-Negative control
8-Negative control

**B**

| STR Locus/line | LQT healthy-PBMC | LQT healthy-iPSC#1 | LQT healthy-iPSC#2 |
|---|---|---|---|
| D3S1358 | 16/17 | 16/17 | 16/17 |
| vWA | 17/19 | 17/19 | 17/19 |
| D16S539 | 9/12 | 9/12 | 9/12 |
| AMEL | X/Y | X/Y | X/Y |
| D8S1179 | 13/15 | 13/15 | 13/15 |
| D21S11 | 29/30 | 29/30 | 29/30 |
| D18S51 | 18/19 | 18/19 | 18/19 |
| D19S433 | 13/14 | 13/14 | 13/14 |
| TH01 | 8/9.3 | 8/9.3 | 8/9.3 |
| FGA | 22/23 | 22/23 | 22/23 |
| D5S818 | 12/13 | 12/13 | 12/13 |
| D13S317 | 9/11 | 9/11 | 9/11 |
| D7S820 | 10.1/11 | 10.1/11 | 10.1/11 |
| SE33 | 17/21.2 | 17/21.2 | 17/21.2 |
| D10S1248 | 14/16 | 14/16 | 14/16 |
| D1S1656 | 14/17.3 | 14/17.3 | 14/17.3 |
| D2S1338 | 17/24 | 17/24 | 17/24 |
| CSF1PO | 12/12 | 12/12 | 12/12 |
| TPOX | 8/8 | 8/8 | 8/8 |
| Yindel | 1/1 | 1/1 | 1/1 |
| DYS391 | 9/9 | 9/9 | 9/9 |
| D2S441 | 10/10 | 10/10 | 10/10 |
| D22S1045 | 11/11 | 11/11 | 11/11 |
| D12S391 | 18/18 | 18/18 | 18/18 |

| STR Locus/line | LQT-PBMC | LQT-iPSC#1 | LQT-iPSC#2 |
|---|---|---|---|
| D3S1358 | 15/17 | 15/17 | 15/17 |
| vWA | 17/19 | 17/19 | 17/19 |
| D16S539 | 9/11 | 9/11 | 9/11 |
| AMEL | X/X | X/X | X/X |
| D8S1179 | 15/15 | 15/15 | 15/15 |
| D21S11 | 30/32.2 | 30/32.2 | 30/32.2 |
| D18S51 | 16/18 | 16/18 | 16/18 |
| D19S433 | 13/16.2 | 13/16.2 | 13/16.2 |
| TH01 | 8/9.3 | 8/9.3 | 8/9.3 |
| FGA | 21/24 | 21/24 | 21/24 |
| D5S818 | 11/13 | 11/13 | 11/13 |
| D13S317 | 9/11 | 9/11 | 9/11 |
| D7S820 | 10/12 | 10/12 | 10/12 |
| SE33 | 17/21.2 | 17/21.2 | 17/21.2 |
| D10S1248 | 13/14 | 13/14 | 13/14 |
| D1S1656 | 14/14 | 14/14 | 14/14 |
| D2S1338 | 17/24 | 17/24 | 17/24 |
| CSF1PO | 12/12 | 12/12 | 12/12 |
| TPOX | 8/8 | 8/8 | 8/8 |
| Yindel | | | |
| DYS391 | 10/10 | 10/10 | 10/10 |
| D2S441 | 10/11 | 10/11 | 10/11 |
| D22S1045 | 11/11 | 11/11 | 11/11 |
| D12S391 | 18/18 | 18/18 | 18/18 |

**C**

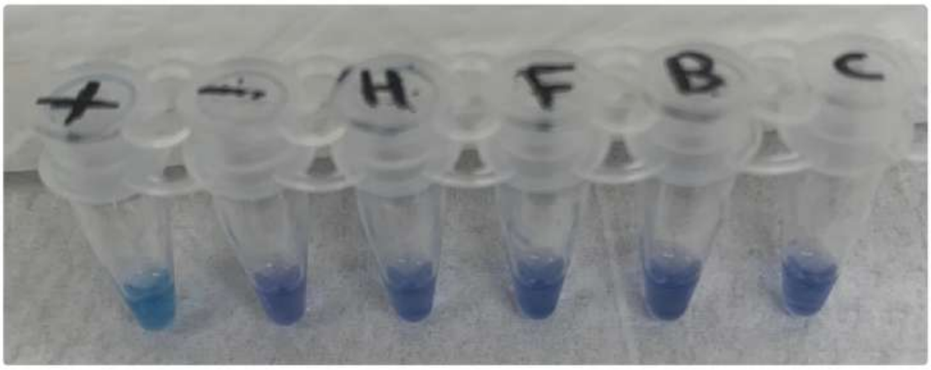


+: Positive control
-: Negaitive control
H: LQT iPSC#1
F: LQT iPSC#2
B: LQT healthy iPSC#1
C: LQT healthy iPSC#2
